# Prediction of breast cancer proteins using molecular descriptors and artificial neural networks: a focus on cancer immunotherapy proteins, metastasis driver proteins, and RNA-binding proteins

**DOI:** 10.1101/840108

**Authors:** Andrés López-Cortés, Alejandro Cabrera-Andrade, José M. Vázquez-Naya, Alejandro Pazos, Humberto Gonzáles-Díaz, César Paz-y-Miño, Santiago Guerrero, Yunierkis Pérez-Castillo, Eduardo Tejera, Cristian R. Munteanu

**Author notes:** These authors contributed equally to the study (Andrés López-Cortés, Alejandro Cabrera-Andrade). **e-mail of authors:** Andrés López-Cortés, Alejandro Cabrera-Andrade, José M. Vázquez-Naya, Alejandro Pazos, Humberto Gonzáles-Díaz, César Paz-y-Miño, Santiago Guerrero, Yunierkis Pérez-Castillo, Eduardo Tejera, Cristian R. Munteanu. **Correspondence:** Andrés López-Cortés, MSc., Centro de Investigación Genética y Genómica, Facultad de Ciencias de la Salud Eugenio Espejo, Universidad UTE, Mariscal Sucre Avenue, Quito 170129, Ecuador.

## Abstract

**Background:** Breast cancer (BC) is a heterogeneous disease characterized by an intricate interplay between different biological aspects such as ethnicity, genomic alterations, gene expression deregulation, hormone disruption, signaling pathway alterations and environmental determinants. Due to the complexity of BC, the prediction of proteins involved in this disease is a trending topic in drug design.

**Methods:** This work is proposing accurate prediction classifier for BC proteins using six sets of protein sequence descriptors and 13 machine learning methods. After using a univariate feature selection for the mix of five descriptor families, the best classifier was obtained using multilayer perceptron method (artificial neural network) and 300 features.

**Results:** The performance of the model is demonstrated by the area under the receiver operating characteristics (AUROC) of 0.980 ± 0.0037 and accuracy of 0.936 ± 0.0056 (3-fold cross-validation). Regarding the prediction of 4504 cancer-associated proteins using this model, the best ranked cancer immunotherapy proteins related to BC were RPS27, SUPT4H1, CLPSL2, POLR2K, RPL38, AKT3, CDK3, RPS20, RASL11A and UBTD1; the best ranked metastasis driver proteins related to BC were S100A9, DDA1, TXN, PRNP, RPS27, S100A14, S100A7, MAPK1, AGR3 and NDUFA13; and the best ranked RNA-binding proteins related to BC were S100A9, TXN, RPS27L, RPS27, RPS27A, RPL38, MRPL54, PPAN, RPS20 and CSRP1.

**Conclusions:** This powerful model predicts several BC-related proteins which should be deeply studied to find new biomarkers and better therapeutic targets. The script and the results are available as a free repository at https://github.com/muntisa/neural-networks-for-breast-cancer-proteins.

## INTRODUCTION

The intricate interplay between several biological aspects such as environmental determinants, gene expression deregulation, genetic alterations, signaling pathway alterations and ethnicity causes the development of breast cancer (BC) heterogeneous disease [1–4]. Over the last years, multi-omics studies, pharmacogenomics treatments and precision medicine strategies have evolved favorably; however, there are still biases such as the significant inclusion of minority populations in cancer research [5–9]. Nowadays, BC is the most commonly diagnosed cancer (2,088,849; 24% cases), and the leading cause of cancer-related deaths among women (626,679; 15% cases) worldwide [10].

In our previous study, López-Cortés *et al.* developed the OncoOmics strategy to reveal essential genes in BC [11]. This strategy was a compendium of approaches that analyzed genetic alterations, protein expression, protein-protein interactions (PPi) and dependency maps of BC genes/biomarkers using relevant databases such as the Pan-Cancer Atlas project [5, 12–14], The Cancer Genome Atlas (TCGA) [15], The Human Protein Atlas (HPA) [16–18], the DepMap project [19–21], and the OncoPPi network [22].

Gene sets were taken from the Consensus Strategy (CS) [23], the Pan-Cancer Atlas, [5, 12–14], the Pharmacogenomics Knowledgebase (PharmGKB) [24, 25], and the Cancer Genome Interpreter (CGI) [26]. The CS, developed by López-Cortés *et al.* and Tejera *et al.*, was proved to be highly efficient in the recognition of genes associated with BC pathogenesis [23, 27]. The Pan-Cancer Atlas reveals how genetic alterations such as protein expression, copy number variants (CNV), mRNA expression, fusion genes and putative mutations collaborate in BC progression [5, 12, 13, 28–33]. PharmGKB is a comprehensive resource that collects the precise guidelines for the application of pharmacogenomics in clinical practice [24, 25]. Lastly, the CGI flags genomic biomarkers of drug response with different levels of clinical relevance [26].

The BC essential genes were rationally filtered to 144 [34], 5 (3.5%) of them were significant in all OncoOmics approaches: RAC1, AKT1, CCND1, PIK3CA and ERBB2, and 15 (10%) were significant in three OncoOmics approaches: CDH1, MAPK14, TP53, MAPK1, SRC, RAC3, PLCG1, GRB2, MED1, TOP2A, GATA3, BCL2, CTNNB1, EGFR and CDK2 [11]. On the other hand, g:Profiler let us know the enrichment map of the 144 essential genes in BC [35]. The most significant gene ontologies (GO) related to biological process and molecular function were the positive regulation of macromolecule metabolic process and the phosphatidylinositol 3-kinase activity, respectively. The most significant term according to the Human Phenotype Ontology was breast carcinoma [36]. Subsequently, the most relevant network interactions of the GO: biological process and the Reactome pathways were related to the immune system [37], tyrosine kinase [38], cell cycle [39], DNA repair [40], and RNA-binding proteins [41].

According to the Open Targets Platform [42], the largest number of drugs that are being analyzed in clinical trials to treat BC with a direct focus on the essential genes were small molecules that correspond most likely to tyrosine kinases. Hence, the essential proteins with signaling function are the best targets for drugs in order to modify any biological activity.

Due to the fact that the experimental methods are expensive and time-consuming, the theoretical methods could offer the practical solution for this screening. Therefore, classification models that link the protein structure to the signaling activity could be obtained using machine learning (ML) techniques, encoding molecular information into invariant molecular descriptors based on molecular graph topology, 3D protein conformation, protein sequence and physical-chemical properties of the amino acids. The classification model represents a quantitative structure-activity relationship (QSAR) between the protein structure and biological function [43]. Different classification models have been published for prediction of protein activities: anti-oxidant [44], lectins [45], signaling [46], anti-angiogenic [47], anti-cancer [48], and enzyme class [49]. Vilar *et al.* [50] developed a QSAR model for alignment-free prediction of BC cancer biomarkers using linear discriminant analysis method, electrostatic potentials of protein pseudofolding HP-lattice networks as features, and 122 proteins related to BC and a control group of 200 proteins with classifications above 80%. Our group [51], proposed an improved multi-target classification model for human breast and colon cancer-related proteins by using a similar molecular graph theory for descriptors: star graph topological indices. The accuracies of the models were 90.0% for a linear forward stepwise model. Both models presented linear relationships between graph-based protein sequence descriptors and BC, and unbalanced datasets. Thus, the aim of this study was to obtain an effective machine learning classification model to predict BC-related proteins screening cancer immunotherapy proteins (CIPs), metastasis driver proteins (MDPs) and RNA-binding proteins (RBPs), using non-graph protein sequence descriptors and additional non-linear ML techniques.

## METHODS

Fig. 1 presents the general flow chart of the methodology to obtain a classifier for BC proteins. In the first step, we constructed a database with BC proteins and non-cancer proteins. In the second step, five families of Rcpi (R package) [52] molecular descriptors have been used: 20 amino acid composition (AC), 400 di-amino acid composition (DC), 8000 tri-amino acid composition (TC), 80 amphiphilic pseudo-amino acid composition (APAAC), and 240 normalized Moreau-Broto autocorrelation (MB). The six sets of descriptors were constructed by mixing all the five-descriptor families, resulting 8708 total descriptors (Mix).

**Fig. 1.**
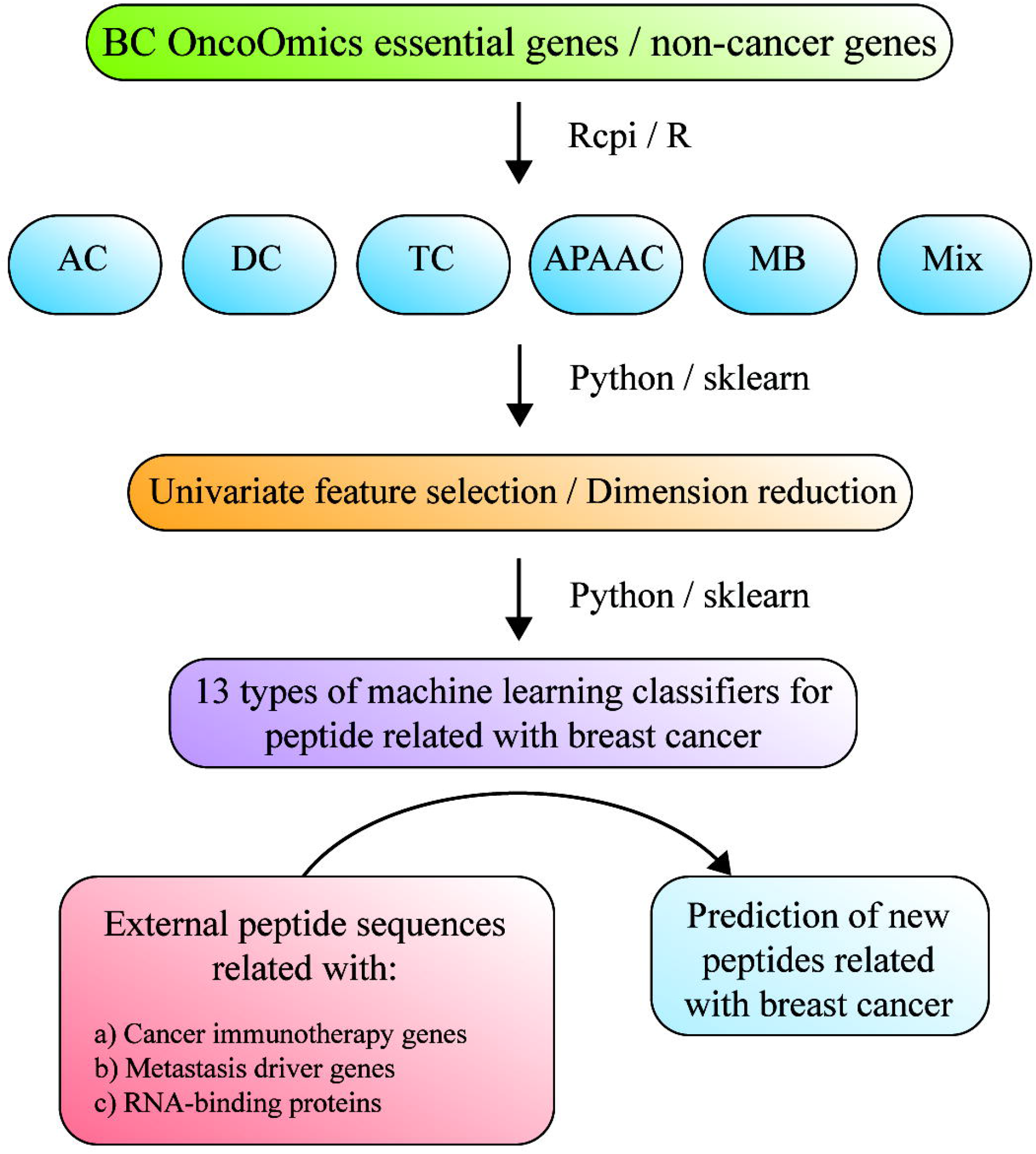
Flow chart of methodology for breast cancer protein prediction.

Jupyter notebooks with python/sklearn [53] were used to test 13 types of ML classifiers for each set of descriptor, without feature selection, with univariate feature selection or using principal component analysis (PCA) [54]. The classifiers were Gaussian Naive Bayes (NB) [55], k-nearest neighbors algorithm (KNeighborsClassifier) [56], linear discriminant analysis (LinearDiscriminantAnalysis) [57], support vector machine (SVM linear and SVM non-linear based on radial basis functions (RBF), support vector classification (SVC kernel = linear) and SVC kernel = RBF [58], logistics regression (LogisticRegression) [59], multilayer perceptron / neural network (MLPClassifier) with 20 neurons in one hidden layer [60], decision tree (DecisionTreeClassifier) [61], random forest (RandomForestClassifier) [62], XGBoost (XGB) is an optimized and distributed gradient boosting library (XGBClassifier) [63], Gradient Boosting for classification (GradientBoostingClassifier) [64], AdaBoost classifier (AdaBoostClassifier) [65], and Bagging classifier (BaggingClassifier) [66]. The feature selection methods was univariate filter as SelectKBest(chi2, k) and the dimension reduction technique was PCA [54].

NB is based on Bayes’ theorem and considers all the features are independent [55]. k-nearest neighbors algorithm (KNN) assigns an unclassified sample using the nearest of k samples in the training set [56]. Linear discriminant analysis (LDA) is a basic linear classifier [57]. Support vector machine (SVM linear and SVM) is using a higher dimensionality space to map the input features [58]. For nonlinear problems, SVM uses Gaussian radial basis (RBF) as nonlinear kernels.

Logistics regression (LG) is another linear classifier that is able to calculate probability of a binary response using weights [59]. Multilayer perceptron (MLP) represents a basic neural network with one hidden layer and with an ability to combine linear and nonlinear functions inside artificial neurons [60]. Decision tree (DT) (DecisionTreeClassifier) represents a tree-type structure of decision rules obtained from the inputs [61]. Random forest (RF) is an ensemble method that combines parallel decision trees [62]. XGBoost (XGB) uses sequential weak trees to improve the classification performance [63]. Gradient Boosting for classification (GB) is a basis boost method using sequential weak classifiers [64]. AdaBoost classifier (AdaB) is mixing different classifiers: it starts the fitting with a classifier based on the original dataset and adds additional copies of the original classifier with adjusted weights for the incorrectly classified instances [65]. Bagging classifier (Bagging) is a modified version of AdaB: the additional classifiers are based on subsets of the original dataset [66].

The ML prediction model was constructed from two protein sets; the positive set (BC-related proteins) was made up of 143 proteins according to López-Cortés *et al.* [11]. On the other hand, the negative protein set was made up of 233 non-cancer proteins (http://www.galseq.com/oncoscore.html) [67]. Tables S1 and S2 detail the sets and FASTA sequences of BC-related proteins and non-cancer proteins, respectively.

Three lists of cancer-related proteins were scanned with the final ML prediction model: 1232 CIPs were taken from Patel *et al.* [37], 1903 MDPs were taken from the Human Cancer Metastasis Database (HCMDB) (http://hcmdb.i-sanger.com/index) [68], and 1369 RBPs were taken from Hentze *et al.*, [41, 68] (Tables S3-S5).

After the calculation of amino acid composition descriptors, the datasets contained 376 proteins. The BC class was labeled with 1 and non-cancer class with 0. Several preprocessing was done before any calculation: elimination of dubled examples, elimination of data with NA values, and elimination of features with zero variance. All feature values were normalized to values between 0 and 1 using MinMax() scaler. A SMOTE filter was used [69] to balance the dataset. The performance of the models used Area Under the Receiver Operating Characteristics (AUROC) [70] metrics and 3-fold cross-validation (CV) method. The best model to be used for predictions was chosen using criteria such as mean AUROC, standard deviation (SD) of AUROC, and the number of features. All the results can be reproduced by using the scripts and datasets at https://github.com/muntisa/neural-networks-for-breast-cancer-proteins.

After the screening of the three cancer-related protein sets through the ML model, the proteins related to breast cancer were analyzed by using g:Profiler (https://biit.cs.ut.ee/gprofiler/) in order to obtain significant annotations (FDR < 0.001) related to GO terms and pathways [35, 41, 68].

Finally, a complementary analysis was done using data of genetic alterations (putative mutations, fusion genes, mRNA expression and copy number variants) according to the Pan-Cancer Atlas [5, 12–14]. We compared the amount of genetic alterations from 981 BC patients among the BC essential proteins, non-cancer proteins, top 100 CIPs related to BC, top 100 MDPs related to BC and top 100 RBPs related to BC. All genomics data can be visualized in the Genomic Data Commons of the National Cancer Institute and in the cBioPortal (https://www.cbioportal.org/) [71, 72].

## RESULTS AND DISCUSSION

The current work proposes better classification models to predict new breast cancer proteins using six sets of protein sequence descriptors calculated with Rcpi: AC, DC, TC, APAAC, MB and Mix. 507 ML classifiers were built using python, 13 types of ML classifiers (NB, KNN, LDA, SVM linear, SVM, LR, MLP, DT, RF, XGB, GB, AdaB and Bagging), univariate filter as feature selection method, and PCA transformation of features. All the models used as performance metrics AUROC (mean values using 3-fold CV).

For the first models, we used the pool of features for the six sets of descriptors without any feature selection or dimension reduction with 12 ML methods (Fig. 2). We can observe that with a big number of descriptors in TC and Mix (over 8000), it is possible to obtain mean AUROC values greater than 0.9 with SVM linear, LR, and MLP. Even with 20 AC descriptors and XGB it is possible to obtain a mean AUROC of 0.857. But we tried to improve this performance and we applied univariate feature selection or PCA dimension reduction to reduce the number of inputs to a maximum of 300 features (due to the small number of instances).

**Fig. 2.**
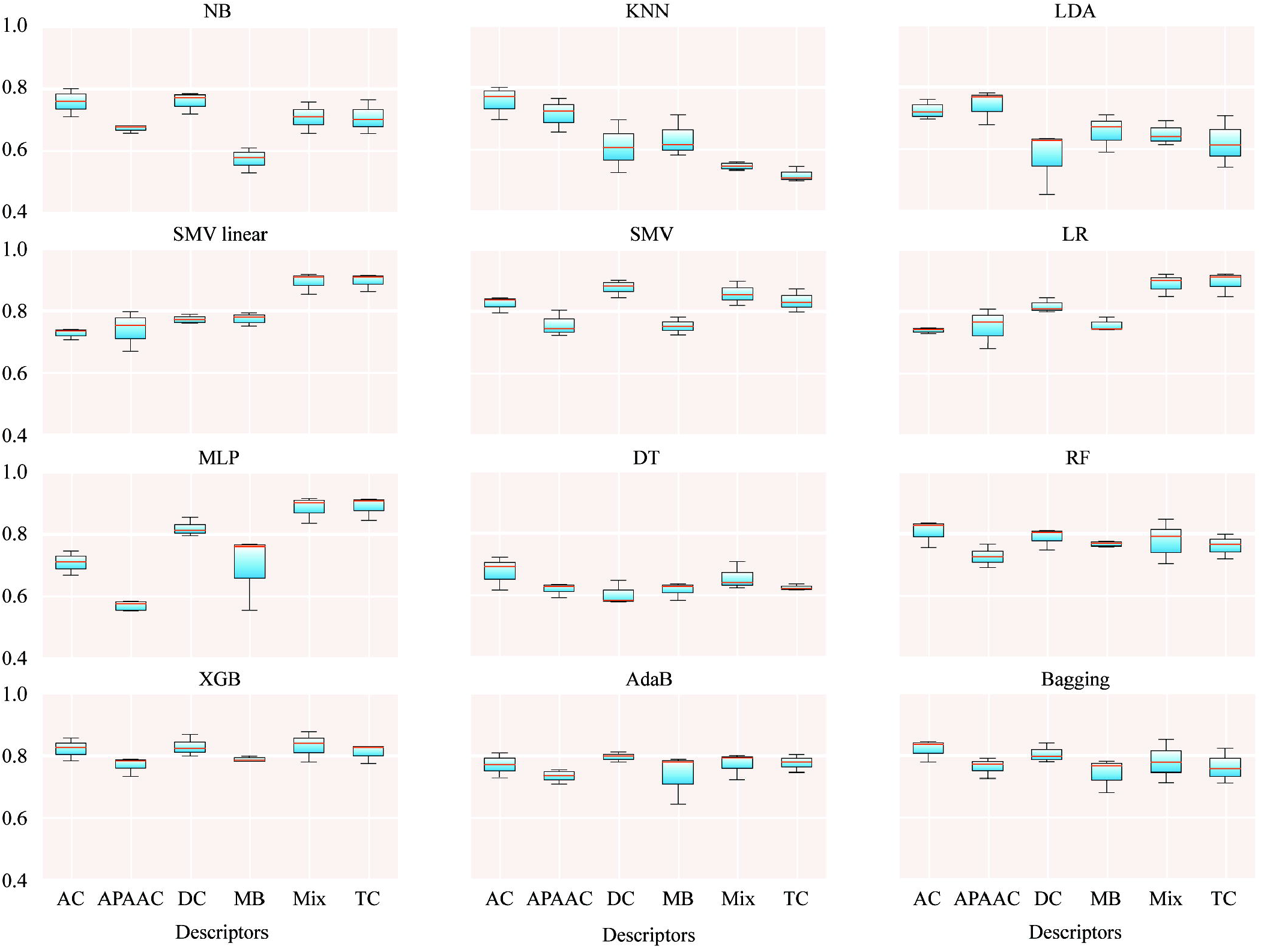
Mean AUROC of classifiers for breast cancer proteins using all features.

Therefore, we selected models based on 20, 100, 200, and 300 features (see 1-ML-BreastCancerPeptides.ipynb). Fig. 3 presents mean AUROC values for classifiers based on only 20 features: AC, DS-Best20, DC-PCA20, TC-Best20, TC-PCA20, APAAC-Best20, APAAC-PCA20, MB-Best20, MB-PCA20, Mix-Best20 and Mix-PCA20 (Best = univariate filter, PCA = feature transformation). DS-Best20 with only 20 di-amino acid composition descriptors and Mix-Best20 with a mixture of descriptors are able to offer mean AUROC values over 0.84 with non-linear SVM, XGB and GB. Additional results could be found in Table S6.

**Fig. 3.**
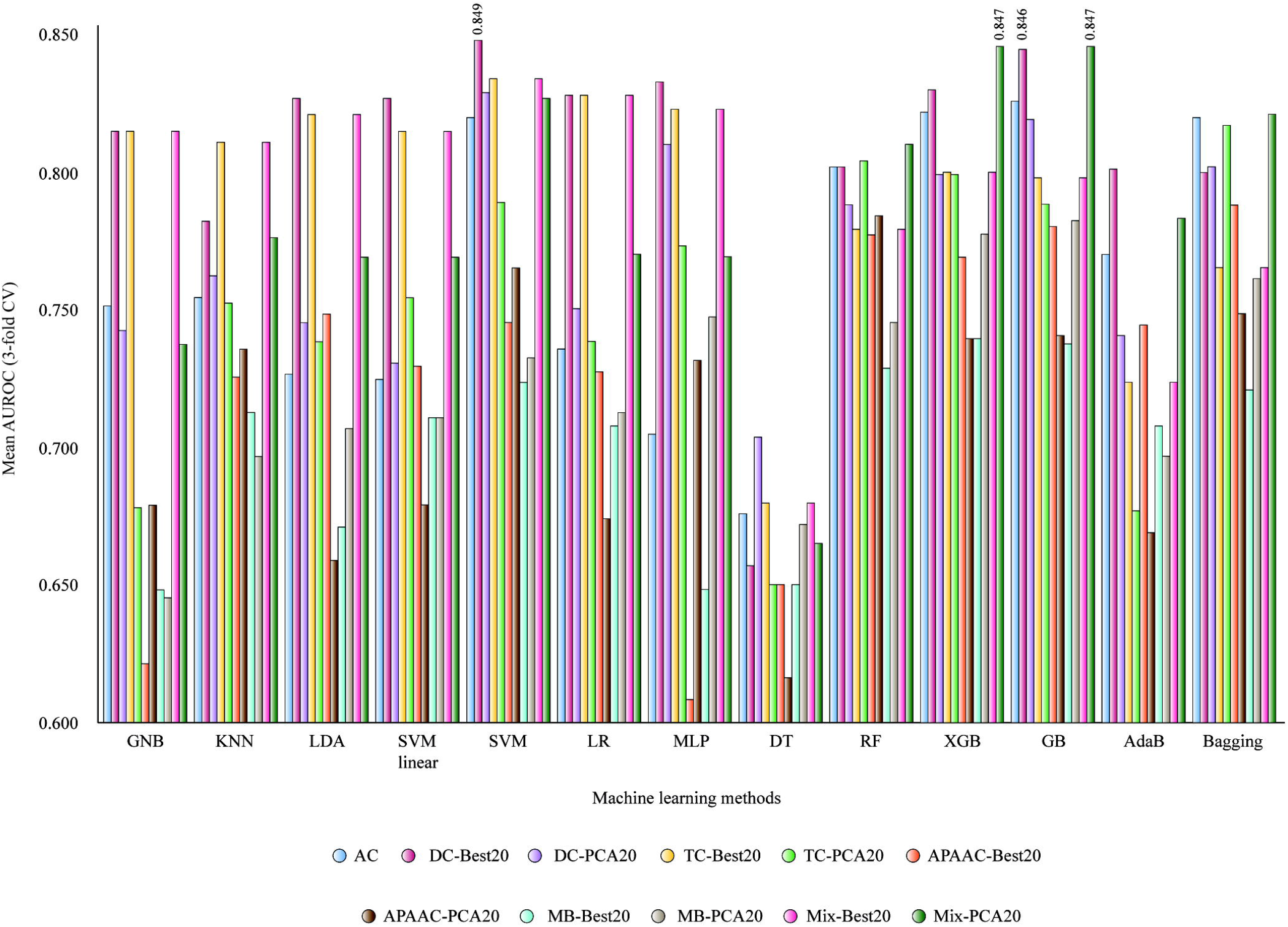
Mean AUROC values for classifiers obtained with 20 selected features (3-fold CV).

If the number of features increased to 100 (5 times from 20), better AUROC values are obtained in Fig. 4: DC-Best100, DC-PCA100, TC-Best100, TC-PCA100, MB-Best100, MB-PCA100, Mix-Best100, and Mix-PCA100. Two sets of descriptors with four ML methods are able to provide mean AUROC values greater than 0.9: TC-Best100 and Mix-Best100 with SVM linear, non-linear SVM, LR and MLP. Thus, LR and TC-Best100 (100 descriptors of tri-amino acid composition) generate a classifier with mean AUROC of 0.917. The increasing of AUROC values is important from 20 to 100 best descriptors. In the next step, the number of selected descriptors was increased to 200. The PCA transformed sets using the same number of components, as the selected features are not able to provide similar classification performance.

**Fig. 4.**
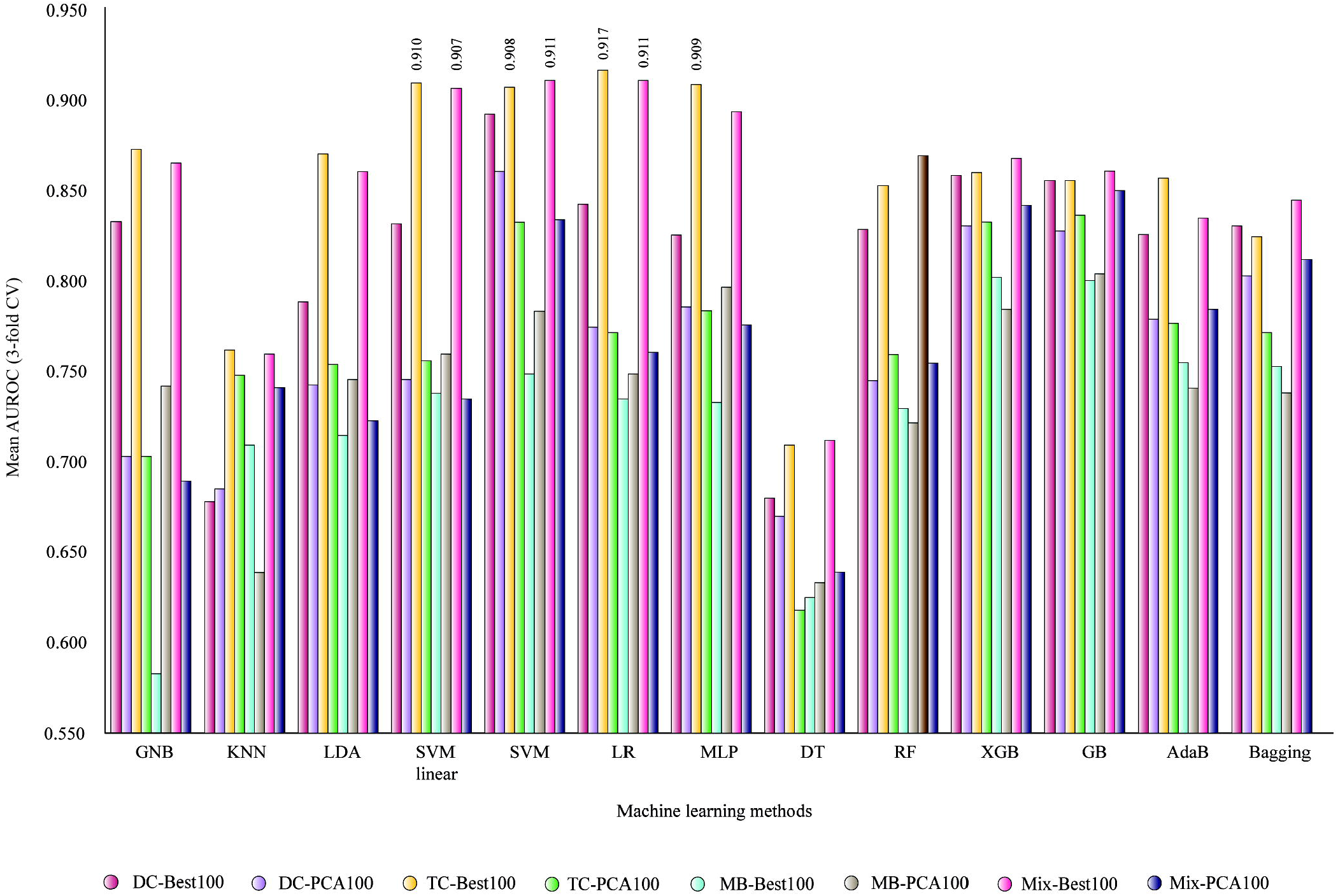
Mean AUROC for classifiers based on 100 selected features (3-fold CV).

Fig. 5 presents the AUROC values for classifiers based on 200 selected features (a double number of inputs from 100): DC-Best200, DC-PCA200, TC-Best200, TC-PCA200, MB-Best200, MB-PCA200, Mix-Best200, and Mix-PCA200. We can observe that the same TC and Mix-based sets are providing mean AUROC values between 0.90 and 0.95 with five ML methods: NB, SVM linear, LR, MLP, and RF. The maximum mean AUROC value was 0.950 using TC-Best200 and the simple linear LR method.

**Fig. 5.**
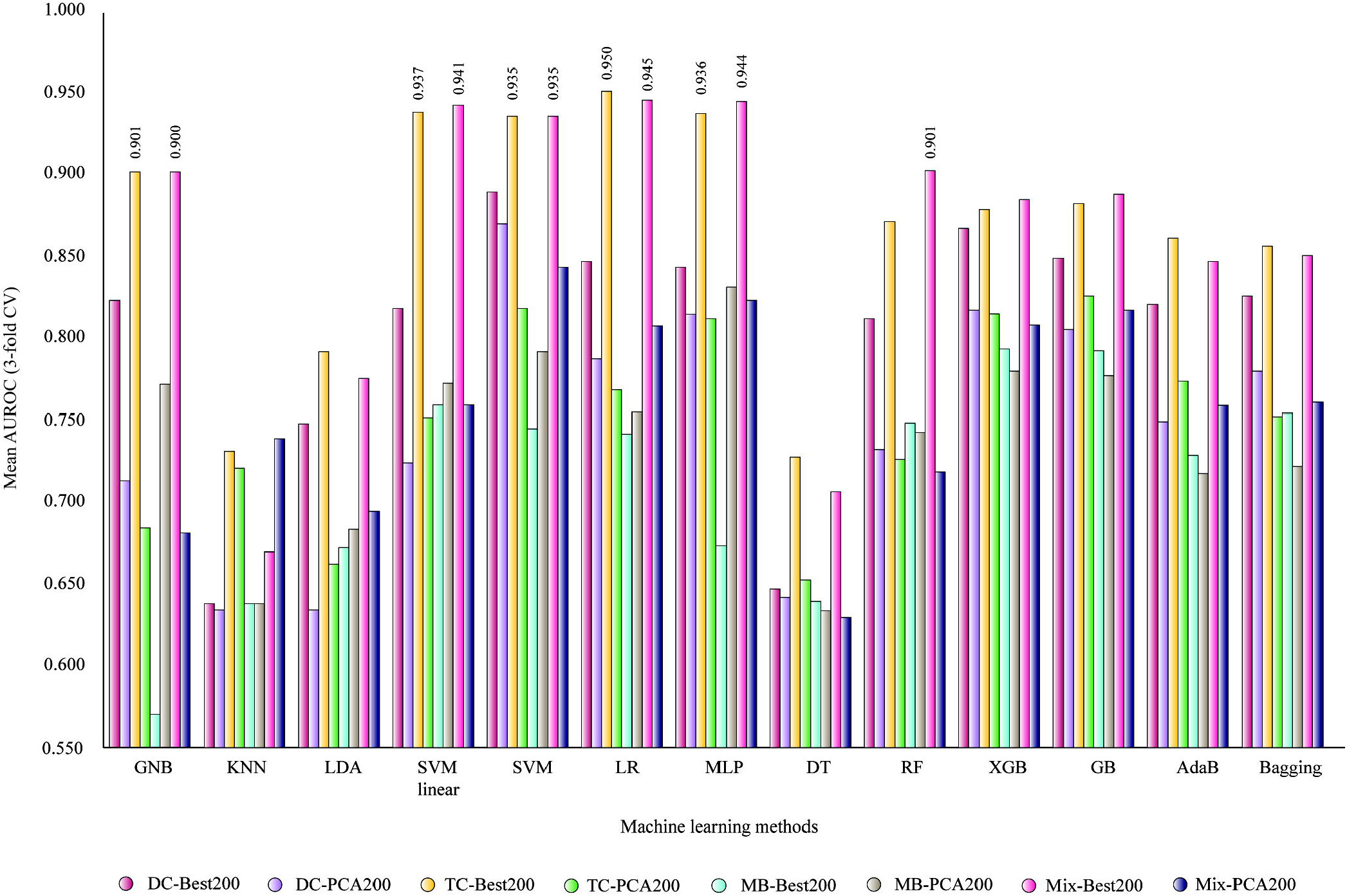
Mean AUROC of classifiers based on 200 selected features (3-fold CV).

In Fig. 6 are presented the AUROC values for classifiers based on 300 selected features: DC-Best300, DC-PCA300, TC-Best300, TC-PCA300, Mix-Best300, and Mix-PCA300. With 300 features, it is possible to provide more accurate classifier for BC proteins. The same TC and Mix subsets can generate classifier with mean AUROC from 0.963 to 0.980 using SVM linear, SVM, LR and MLP.

**Fig. 6.**
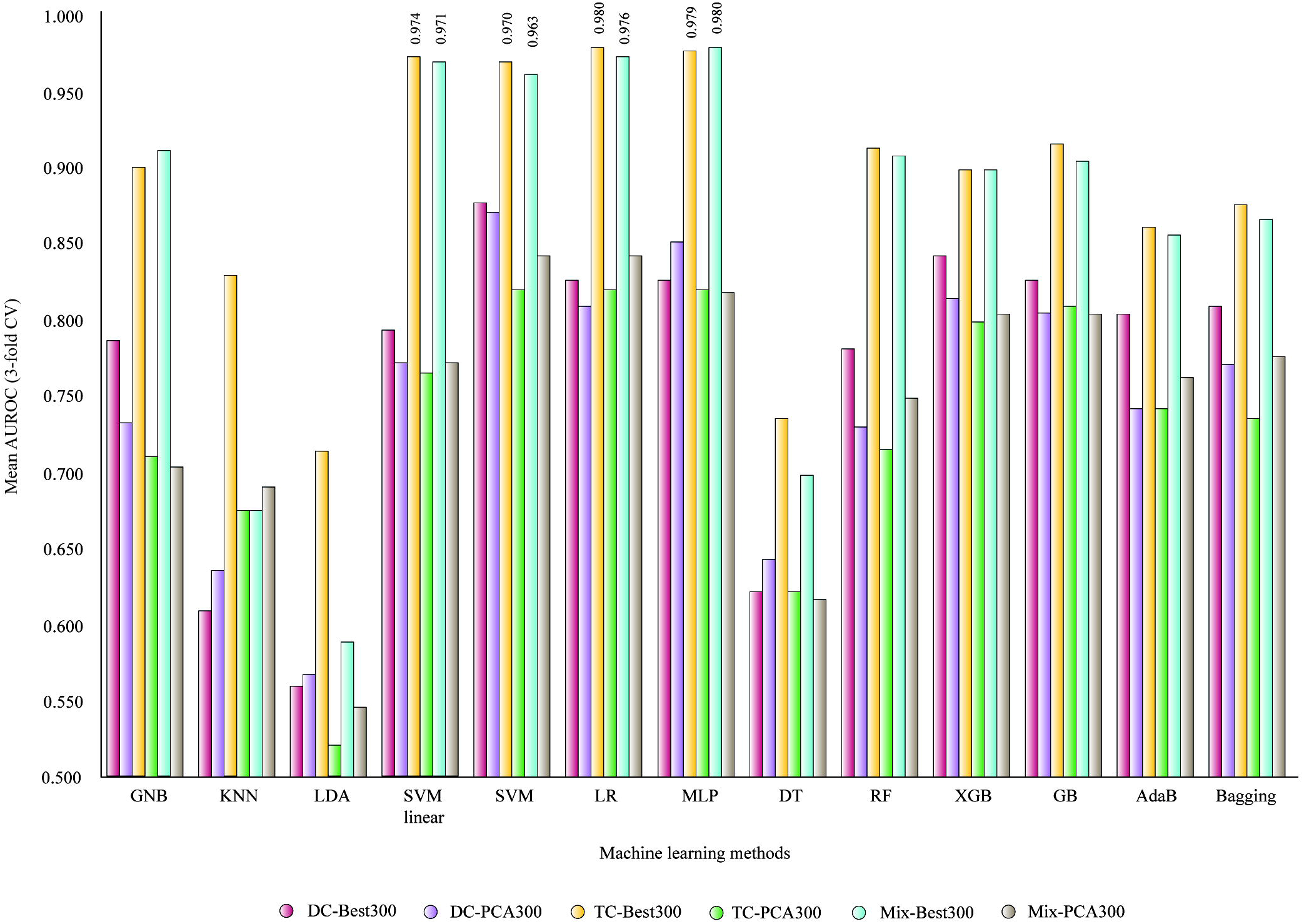
Mean AUROC of classifiers based on 300 selected features (3-fold CV).

The best AUROC of 0.980 ± 0.0037 was obtained with MLP and Mix-Best300. The same AUROC value was generated by TC-Best300 and LR but with a double SD of 0.0077. In the best model with the mixed descriptors, between the 300 descriptors, seven DC (LG, QI, NK, EM, QM, MM and EY) and two APAAC descriptors (Pc1.N and Pc1.M) were selected for BC function. The rest are TC descriptors without any MP descriptor selected (see Table S7). The accuracy of the best model was 0.936 ± 0.0056.

4504 external proteins were transformed in the molecular descriptors of the best model and were used to predict its breast cancer activity (see 2-Predictions-BreastCancerPeptides.ipynb): 1232 cancer immunotherapy proteins, 1903 metastasis driver proteins, and 1369 RNA-binding proteins. Thus, all these proteins were transformed into 300 selected descriptors of Mix-300 set and used them with the saved MLP classifier. The results are presented in Tables S3-S5.

After obtaining the list of proteins predicted as BC-related, we performed a functional enrichment analysis through g:Profiler to interpret the GO of these proteins [73]. Figs. 7A and 7B show the enrichment map of the 608 cancer immunotherapy proteins related to BC. g:Profiler searches for a collection of proteins representing pathways, GO terms, and disease phenotypes [73]. The most significant false discovery rate (FDR < 0.001) GO: molecular functions were RNA-binding, structural constituent of ribosome and nucleic acid binding; the most significant GO: biological processes were mRNA metabolic process, protein targeting to estrogen receptor and SRP-dependent cotranslational protein targeting to membrane; the most significant Kyoto Encyclopedia of Genes and Genomes (KEGG) pathway was ribosome [74]; the most significant Reactome pathways were Cap-dependent translation initiation, eukaryotic translation initiation and formation of a pool of free 40S subunits [74, 75]; lastly, the most significant WikiPathway was cytoplasmic ribosomal proteins [76]. The g:Profiler terms are fully detailed in Table S8.

**Fig. 7.**
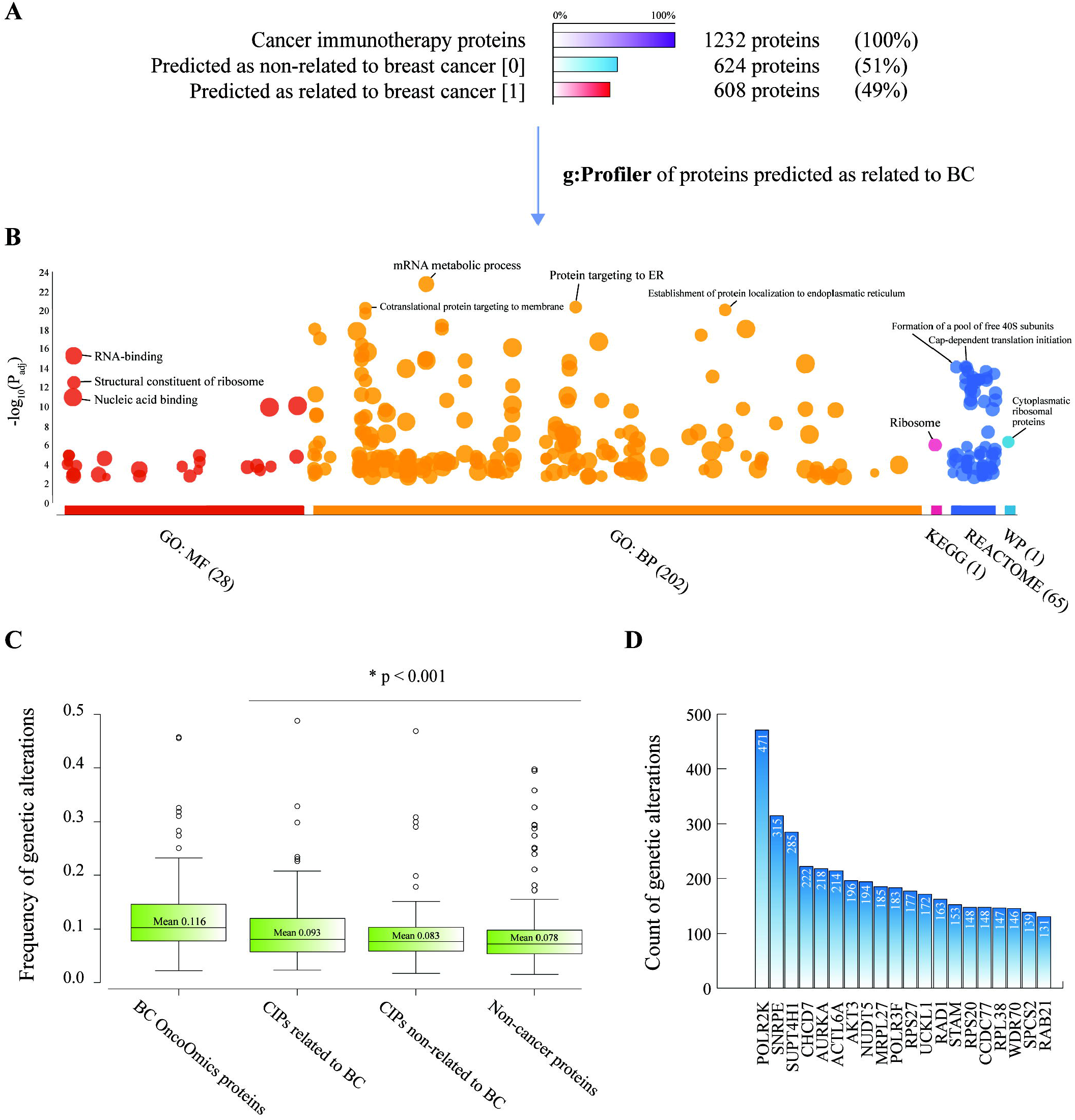
Cancer immunotherapy proteins. A) Percentage of CIPs related to BC. B) g:Profiler enrichment of CIPs related to BC. C) Frequency of genetic alterations per sample between BC proteins, CIPs related to BC, CIPs non-related to BC and non-cancer proteins. D) CIPs with the highest number of genetic alterations according to 981 individuals from the Pan-Cancer Atlas.

Cancer immunotherapy is revolutionizing oncology. Given the success in achieving long-term durable responses in numerous advanced and metastatic solid cancers, cancer immunotherapy sparked tremendous interest and research activities in basic, translational and clinical science [77]. The 10 CIPs best related to BC according to our ML predictions were RPS27, SUPT4H1, CLPSL2, POLR2K, RPL38, AKT3, CDK3, RPS20, RASL11A and UNTD1 (Table S3). For instance, Atsuta *et al.* identified RPS27 as a member of a tumor associated antigen in patients with BC [78].

The Pan-Cancer Atlas is the most sweeping cross-cancer analysis yet undertaken from The Cancer Genome Atlas (TCGA) [12, 71, 72]. Even though, we performed an analysis to compare data of genetic alterations from 981 BC patients [5, 12–14]. Fig. 7C details the frequency of genetic alterations per sample between BC essential proteins (0.116), the top 100 CIPs related to BC (0.093), the top 100 CIPs non-related to BC (0.083) and non-cancer proteins (0.078), being statistically significant the difference of genetic alteration frequencies between CIPs related to BC and non-cancer proteins after the Mann-Whitney U test (p < 0.001). Additionally, the cancer immunotherapy proteins with the greatest number of genetic alterations were POLR2K, SNRPE, SUPT4H1, CHCD7, AURKA, ACTL6A, AKT3, NUDT5, MRPL27 and POLR3F (Table S9) (Fig. 7D).

Figs. 8A and 8B show the enrichment map of the 971 metastasis driver proteins related to BC. The most significant (FDR < 0.001) GO: molecular functions were signaling receptor binding, protein binding and enzyme binding; the most significant GO: biological processes were positive regulation of cellular process, positive regulation of biological process and response to organic substance [73]; the most significant KEGG pathways were pathways in pancreatic, prostatic, colorectal, endometrial and breast cancer types [74]; the most significant Reactome pathways were signaling by interleukins, cytokine signaling in immune system and interleukin-4 and −12 signaling [74, 75]; lastly, the most significant WikiPathways were VEGFA-VEGFR2 signaling pathway and RAC1/PAK1/p38/MMP2 pathway [76]. The g:Profiler terms are fully detailed in Table S10.

**Fig. 8.**
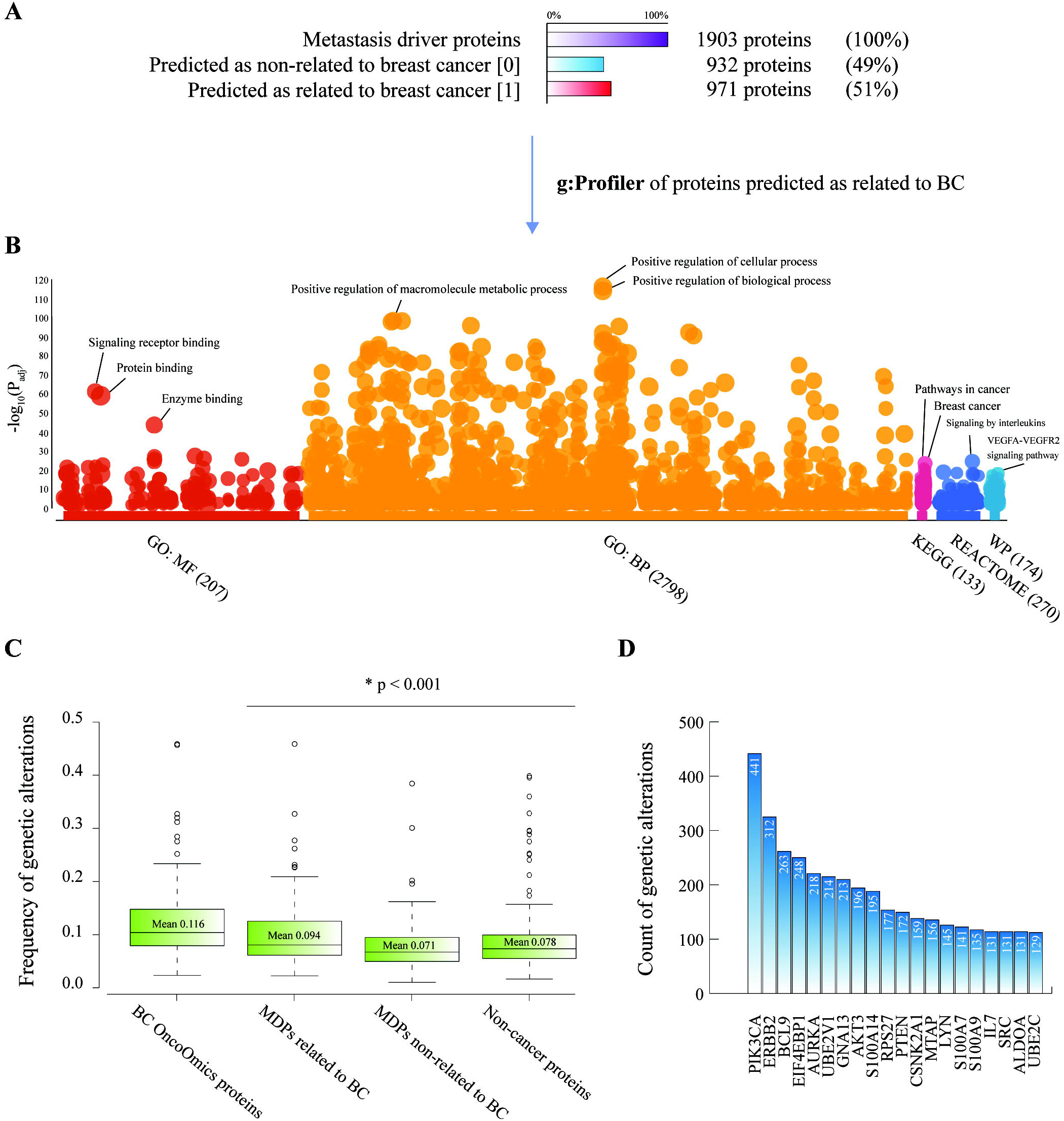
Metastasis driver proteins. A) Percentage of MDPs related to BC. B) g:Profiler enrichment of MDPs related to BC. C) Frequency of genetic alterations per sample between BC proteins, MDPs related to BC, MDPs non-related to BC and non-cancer proteins. D) MDPs with the highest number of genetic alterations according to 981 individuals from the Pan-Cancer Atlas.

Metastasis, often preceded or accompanied by therapeutic resistance, is the most lethal and insidious aspect of cancer. Patients do not die from their primary breast tumor but as a consequence of metastases. Due to tumor evolution and treatment pressure, the genomic alterations in metastatic BC can differ substantially from the primary tumor [79]. To date, the molecular and microenvironmental determinants of metastasis are largely unknown, as is the timing of systemic spread, hindering effective treatment and prevention efforts [13, 68, 80]. Integrated analysis of multi-dimensional transcriptomic data is important to our understanding of cancer metastasis. Moreover, these data would help us identify gene expression signature associated with metastasis in order to choose appropriate treatment strategies [81, 82]. The 10 MDPs best related to BC according to our ML predictions were S100A9, DDA1, TXN, PRNP, RPS27, S100A14, S100A7, MAPK1, AGR3 and NDUFA13 (Table S4). For instance, Bergenfelz *et al.* suggested that S100A9 expressed in negative estrogen receptor and negative progesterone receptor breast cancers induces inflammatory cytokines and is associated with an impaired overall survival [83].

Regarding the Pan-Cancer Atlas project, the boxplots shown in Fig. 8C compare the frequency of genetic alterations per sample between BC proteins (0.116), the top 100 MDPs related to BC (0.094), the top 100 MDPs non-related to BC (0.071) and non-cancer proteins (0.078), being statistically significant the difference of genetic alteration frequencies between MDPs related to BC and non-cancer proteins after the Mann-Whitney U test (p < 0.001). Additionally, the metastasis driver proteins with the greatest number of genetic alterations were PIK3CA, ERBB2, BLC9, EIF4EBP1, AURKA, UBE2V1, GNA13, AKT3, S100A14 and RPS27 (Table S9) (Fig. 8D).

Figs. 9A and 9B show the enrichment map of the 757 RNA-binding proteins related to BC. The most significant (FDR < 0.001) GO: molecular functions were RNA binding, nucleic acid binding and heterocyclic compound binding; the most significant GO: biological processes were RNA processing, mRNA metabolic process and ribonucleoprotein complex biogenesis [73]; the most significant KEGG pathways were spliceosome, ribosome and ribosome biogenesis in eukaryotes [74]; the most significant Reactome pathways were metabolism of RNA, rRNA processing in the nucleus and cytosol and translation [74, 75]; finally, the most significant WikiPathways were mRNA processing and cytoplasmic ribosomal proteins [76]. The g:Profiler terms are fully detailed in Table S11.

**Fig. 9.**
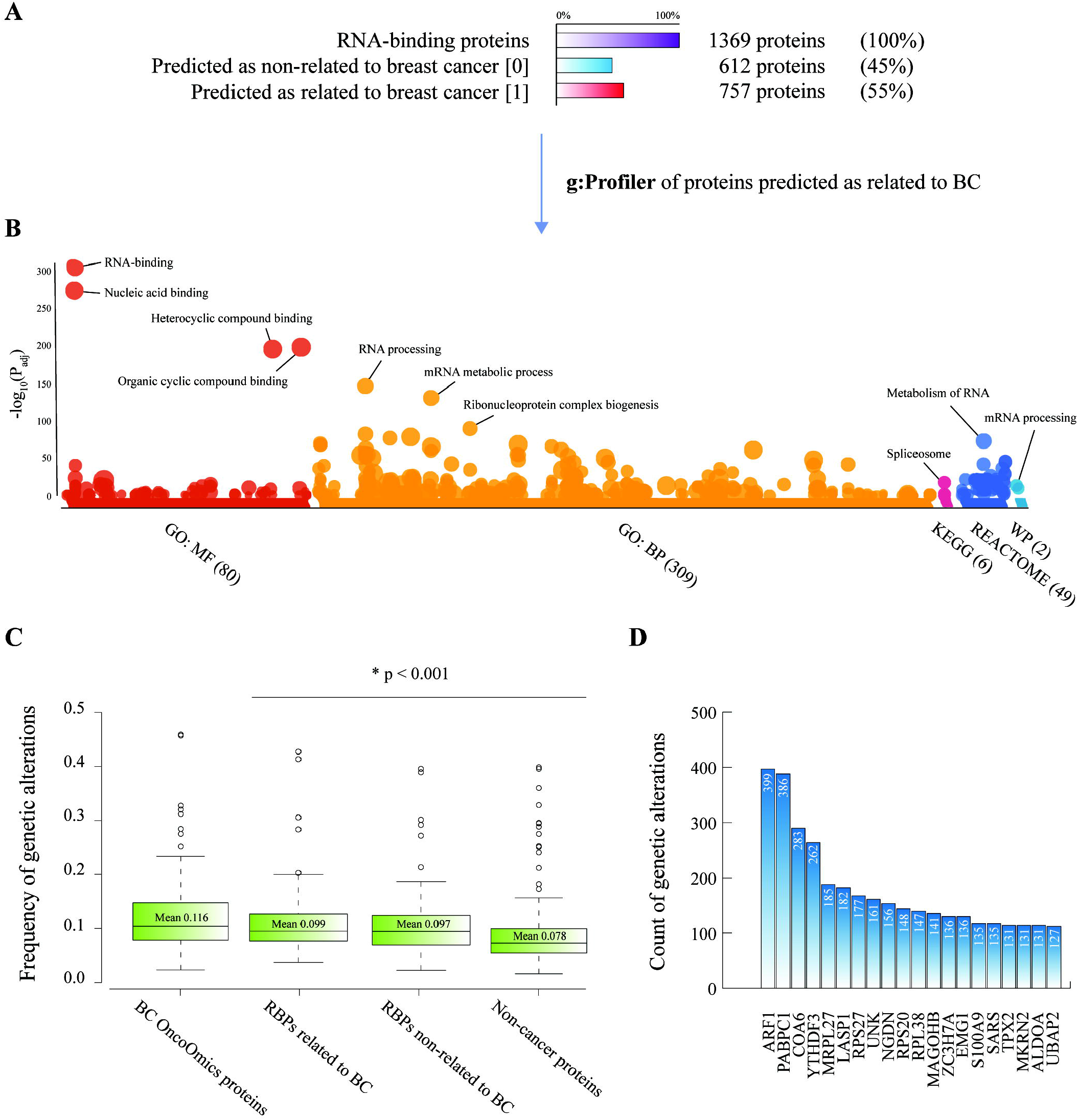
RNA-binding proteins. A) Percentage of RBPs related to BC. B) g:Profiler enrichment of RBPs related to BC. C) Frequency of genetic alterations per sample between BC proteins, RBPs related to BC, RBPs non-related to BC and non-cancer proteins. D) RBPs with the highest number of genetic alterations according to 981 individuals from the Pan-Cancer Atlas.

RNA biology is an under-investigated field of cancer even though pleiotropic changes in the transcriptome are key feature of cancer cell [84]. RBPs are able to control every aspect of RNA metabolism such as translation, splicing, stability, degradation of mRNA, nucleocytoplasmic transport, capping, and polyadenylation [84–87]. RBPs are emerging as critical modulators of BC and the prediction of relation with this complex disease through ML methods provides a better understanding of new genomic targets and biomarkers. The 10 RBPs best related to BC according to our ML predictions were S100A9, TXN, RPS27L, RPS27, RPS27A, RPL38, MRPL54, PPAN, RPS20 and CSRP1 (Table S5). For instance, Rodrigues *et al.* suggested that TXN is overexpressed in BC, and it is related to tumor grade, being a crucial element in redox homeostasis [88].

Regarding the Pan-Cancer Atlas project, Fig. 9C shows the frequency of genetic alterations per sample between BC proteins (0.116), the top 100 RBPs related to BC (0.099), the top 100 RBPs non-related to BC (0.097) and non-cancer proteins (0.078), being statistically significant the difference of genetic alteration frequencies between RBPs related to BC and non-cancer proteins after the Mann-Whitney U test (p < 0.001). Additionally, the RNA-binding proteins with the greatest number of genetic alterations were ARF1, PABPC1, COA6, YTHDF3, MRPL27, LASP1, RPS27, UNK, NGDN and RPS20 (Table S9) (Fig. 9D). In sum, there is an interesting and novel correlation between best predicted BC proteins with machine learning and the amount of pathogenic genetic alterations in CIPs, MDPs and RBPs.

## CONCLUSIONS

The current work proposed better prediction models for breast cancer proteins using as inputs six sets of protein sequence descriptors from Rcpi and 13 machine learning classifiers (with or without feature selection/dimension reduction of features). We choose as the best classifier the MLP classifier. As inputs a mixture of 300 selected molecular descriptors have been used: DC, TC and APAAC. The model has a mean AUROC of 0.980 ± 0.0037 and a mean accuracy of 0.936 ± 0.0056 (3-fold cross-validation). 4504 sequences of proteins related to cancer have been screened for breast cancer relation. The top 10 cancer immunotherapy proteins best related to BC were RPS27, SUPT4H1, CLPSL2, POLR2K, RPL38, AKT3, CDK3, RPS20, RASL11A and UBTD1; the top 10 metastasis driver proteins best related to BC were S100A9, DDA1, TXN, PRNP, RPS27, S100A14, S100A7, MAPK1, AGR3 and NDUFA13; and the top 10 RNA-binding proteins best related to BC were S100A9, TXN, RPS27L, RPS27, RPS27A, RPL38, MRPL54, PPAN, RPS20 and CSRP1. Finally, this powerful model predicts several BC-related proteins that should be deeply studied to find new biomarkers and better therapeutic targets.

## ABBREVIATIONS

BC: breast cancer
PPi: protein-protein interactions
TCGA: The Cancer genome Atlas
HPA: Human Protein Atlas
CS: consensus strategy
CGI: cancer genome interpreter
CNV: copy number variant
GO: gene ontology
ML: machine learning
QSAR: quantitative structure-activity relationship
CIP: cancer immunotherapy protein
MDP: metastasis driver protein
RBP: RNA-binding protein
AC: amino acid composition
DC: di-amino acid composition
TC: tri-amino acid composition
MB: Moreau-Broto
PCA: principal component analysis
NB: Gaussian naive bayes
SVM: support vector machine
RBF: radial basis functions
SVC: support vector classification
KNN: k-nearest neighbors algorithm
LDA: linear discriminant analysis
LG: logistic regression
MLP: multilayer perceptron
DT: decision tree
RF: random forest
GB: gradient boosting
CV: cross-validation
AUROC: area under the receiver operating characteristics
SD: standard deviation
FDR: false discovery rate
PharmGKB: pharmacogenomics knowledgebase
KEGG: Kyoto Encyclopedia of Genes and Genomes
HCMDB: Human Cancer Metastasis Database

## ACKNOWLEDGMENTS

This work was supported by a) the Collaborative Project in Genomic Data Integration (CICLOGEN) PI17/01826 funded by the Carlos III Health Institute from the Spanish National plan for Scientific and Technical Research and Innovation 2013-2016 and the European Regional Development Funds (FEDER) - “A way to build Europe”; b) the General Directorate of Culture, Education and University Management of Xunta de Galicia ED431D 2017/16 and “Drug Discovery Galician Network” Ref. ED431G/01 and the “Galician Network for Colorectal Cancer Research” (Ref. ED431D 2017/23); c) the Spanish Ministry of Economy and Competitiveness for its support through the funding of the unique installation BIOCAI (UNLC08-1E-002, UNLC13-13-3503) and the European Regional Development Funds (FEDER) by the European Union; d) the Consolidation and Structuring of Competitive Research Units - Competitive Reference Groups (ED431C 2018/49), funded by the Ministry of Education, University and Vocational Training of the Xunta de Galicia endowed with EU FEDER funds; e) research grants from Ministry of Economy and Competitiveness, MINECO, Spain (FEDER CTQ2016-74881-P), Basque government (IT1045-16), and kind support of Ikerbasque, Basque Foundation for Science; and, f) Sociedad Latinoamericana de Farmacogenómica y Medicina Personalizada (SOLFAGEM).

## AUTHORS’ CONTRIBUTIONS

ALC, ACA and CRM conceived the subject, the conceptualization of the study and wrote the manuscript. ALC, ACA, JMVN and CRM did data curation and supplementary data. CRM and JMVN built the models using machine learning. AP, HGD, CPyM, SG, YPC and ET gave conceptual advice and valuable scientific input. AP, HGD, CPyM, SM, YPC, ET and CRM supervised the project. ALC and CPyM did funding acquisition. Finally, all authors reviewed the manuscript.

## AVAILABILITY OF DATA AND MATERIALS

All data generated during this study are included in this published article, and the script are available as a free repository at https://github.com/muntisa/neural-networks-for-breast-cancer-proteins.

